# Increased susceptibility to ischemia causes exacerbated response to microinjuries in the cirrhotic liver

**DOI:** 10.1101/2023.07.18.549420

**Authors:** Ben D. Leaker, Mozhdeh Sojoodi, Kenneth K. Tanabe, Yury V. Popov, Joshua Tam, R. Rox Anderson

## Abstract

**Background:** Fractional laser ablation is a technique developed in dermatology to induce remodeling of skin scars by creating a dense pattern of microinjuries. Despite remarkable clinical results, this technique has yet to be tested for scars in other tissues. As a first step towards determining the suitability of this technique, we aimed to (1) characterize the response to microinjuries in the healthy and cirrhotic liver, and (2) determine the underlying cause for any differences in response.

**Methods:** Healthy and cirrhotic rats were treated with a fractional laser then euthanized from 0hr up to 14d after treatment. Differential expression was assessed using RNAseq with a difference-in-differences model. Spatial maps of tissue oxygenation were acquired with hyperspectral imaging and disruptions in blood supply were assessed with tomato lectin perfusion.

**Results:** Healthy rats showed little damage beyond the initial microinjury and healed completely by 7d without scarring. In cirrhotic rats, hepatocytes surrounding microinjury sites died 4-6hr after ablation, resulting in enlarged and heterogeneous zones of cell death. Hepatocytes near blood vessels were spared, particularly near the highly vascularized septa. Gene sets related to ischemia and angiogenesis were enriched at 4hr. Laser-treated regions had reduced oxygen saturation and broadly disrupted perfusion of nodule microvasculature, which matched the zones of cell death.

**Conclusions:** The cirrhotic liver has an exacerbated response to microinjuries and increased susceptibility to ischemia from microvascular damage, likely related to the vascular derangements that occur during cirrhosis development. Modifications to the fractional laser tool, such as using a femtosecond laser or reducing the spot size, may be able to prevent large disruptions of perfusion and enable further development of a laser-induced microinjury treatment for cirrhosis.

## Introduction

Cirrhosis is the liver scarring that occurs as the common final stage of chronic liver diseases, such as viral hepatitis, alcoholic liver disease, and non-alcoholic steatohepatitis. It accounts for over 1 million deaths per year, more than 2% of the global total(1), and ranks 13^th^ in disability-adjusted life years – ahead of malaria, lung cancer, breast cancer, and dementia(2). The only current treatment option is liver transplantation, which is severely limited by short supply of donors and stringent exclusion criteria.

Although our understanding of the disease has improved substantially over the past several decades, little progress has been made in the treatment of cirrhosis. Overall mortality has improved slightly relative to the total population, but this has been attributed to better management of key risk factors, such as alcohol use, rather than substantial improvements in treatment(3). There has been success in developing drugs for the early stages of liver fibrosis, where treatment only needs to stop the injury to allow the innate regenerative abilities of the liver to take over. However, this regenerative capacity is lost once cirrhosis is established. Drug development for fully established cirrhosis has thus far been unsuccessful in part because it is difficult to significantly change tissue architecture with a pharmaceutical approach. In recent years, some of the most promising new research has attempted to use cell therapy to exert a more direct effect(4).

Fibrotic disorders can occur anywhere in the body, but skin scars have the unique property of being easily accessible. Consequently, the development of skin scar therapies has often focused on direct surgical intervention rather than drugs. Examples include z-plasty(5), in which incisions are made along tension lines in the scar, and complete excision of the wound for replacement with an autograft(6). One of the most recent and effective methods is based on microinjury regeneration. While large injuries cause skin to heal by scarring, sufficiently small injuries (nominally <500µm in diameter) induce a wound healing response that leads to healing with normal tissue(7, 8). This property is maintained even in fibrotic skin, and it was found that creating dense patterns of microinjuries in a skin scar using a tool called a fractional laser leads to dramatic improvements and normalization of tissue structure and function. This includes a few clinical reports of the return of complex structures such as hair follicles and sweat glands, which are typically permanently lost in skin scars (7, 9, 10). Microinjury ablation has shown remarkable clinical efficacy, with particularly dramatic results for scars over joints where the stiff, contracted tissue had restricted movement. Tissue stiffness decreases within days of treatment, as shown by substantial improvements in range of motion(11). There is then continued tissue remodeling lasting several months that results in significant improvements in appearance, texture, pliability, pain, and pruritis(7, 11-14). Histology shows neovascularization, reduced density of collagen, and a more organized fibril arrangement (14).

Microinjury ablation is a unique concept that relies on the creation of finely controlled injuries to treat a condition caused by uncontrolled injury. It is unknown how other tissues respond to microinjury ablation. If the treatment can stimulate remodeling of the cirrhotic septa, it may be a powerful new tool in the management of chronic liver disease. It is first necessary to understand how the liver responds to microinjuries. In this study, we aimed to characterize the tissue response to microinjury ablation in the healthy and cirrhotic liver. We then investigated the underlying cause for the differences in response to ablation, with a particular focus on the influence of the liver microvascular architecture.

## Materials & Methods

### Animal model

All animal work was approved by the Massachusetts General Hospital Institutional Animal Care and Use Committee and performed in compliance with the US National Research Council’s Guide for the Care and Use of Laboratory Animals and US Public Health Service’s Policy on Humane Care and Use of Laboratory Animals. Our experiments used male Wistar rats (RRID:RGD_13508588) purchased from Charles River Laboratories. Animals were housed in a controlled environment with food and water ad libitum. For cirrhosis experiments, rats received biweekly injections of thioacetamide at 200mg/kg for 12 weeks followed by a 1 week wash out period.

### Surgery protocol

Animals were anesthetized with isoflurane and prepared for surgery by clipping fur on the abdomen and disinfecting with 10% povidone-iodine. A laparotomy was performed with a 1-2 inch midline incision beginning from the xyphoid. The left lobe of the liver was exposed using sterile swabs. The UltraPulse fractional CO2 laser (Lumenis, Yokneam, Israel) with sterile spacer was brought to the liver surface. The laser ablates a 9×9 array of microinjuries in an approximately 50mm^2^ area (Supplementary Video), with each microinjury receiving two pulses of 15mJ. The majority of animals were treated with the laser in 4 areas, but 2-3 areas were occasionally used depending on the size of the liver. Once bleeding stopped, the muscle and skin were closed with absorbable sutures. Animals were euthanized via cardiac puncture at 0hr, 2hr, 4hr, 6hr, 3d, 7d, and 14d after surgery.

### Histology

Tissue for histological analysis was collected at the time of euthanasia. Samples for hematoxylin & eosin (H&E) and terminal deoxynucleotidyl transferase dUTP nick-end labeling (TUNEL) staining were fixed in 4% formaldehyde, embedded in paraffin blocks, and cut into 5µm sections. TUNEL staining was performed using the DeadEnd Fluorometric TUNEL kit (ProMega, Madison, USA), according to the manufacturer’s instructions. Samples for nitro blue tetrazolium chloride (NBTC; Thermo Fisher Scientific, Waltham, USA) and immunofluorescence staining were embedded in OCT and snap-frozen on dry ice. 10µm (for NBTC) or 5µm (for immunofluorescence) sections were cut in a cryostat at -20°C. NBTC viability staining was performed by incubating the sections with NBTC for 15 min, followed by counterstaining with eosin. For immunostaining, primary antibodies against alpha-smooth muscle actin (SMA, marker of activated hepatic stellate cells; diluted 1:500; Abcam, Cambridge, UK, Cat# ab124964, RRID:AB_11129103), hepatic nuclear factor 4 alpha (HNF4a, hepatocyte marker; 1:100; Abcam Cat# ab41898, RRID:AB_732976), and phosphorylated c-Jun (pcJun, immediate early liver regeneration marker(15); 1:400; Abcam Cat# ab30620, RRID:AB_726902) were used. The sections were then stained with Alexa Fluor 488 or 594 conjugated secondary antibody (Thermo Fisher Scientific) and DAPI (Thermo Fisher Scientific). Slides were imaged with a Nanozoomer 2.0 HT slide scanner (Hamamatsu, Hamamatsu City, Japan).

### RNAseq

Tissue for RNAseq analysis was collected at the 4hr euthanasia timepoint from healthy and cirrhotic rats. Two roughly 25mg samples were collected from each animal – one from an ablated region, and one from an untreated region in the same lobe to be used as a control – and placed in RNA*later* (Thermo Fisher Scientific). Samples were stored at 4°C overnight and then at -20°C until use. RNA extraction was performed using the RNeasy midi kit with proteinase K digestion and on-column DNase digestion according to the manufacturer’s instructions (Qiagen, Hilden, Germany). The samples were then sent to Admera Health for quality control, library preparation, and sequencing. Libraries were prepared using NEBNext Ultra II (New England Biolabs, Ipswich, USA) with poly(A) selection and sequenced on a NovaSeq 6000 (Illumina, San Diego, USA) with 40M 150bp paired-end reads per sample.

Sequencing data quality was assessed with FastQC(16) and trimmed with Cutadapt(17). Transcript abundance was quantified with the Salmon quasi-mapping tool(18). Differential expression analysis was performed using DESeq2 (RRID:SCR_015687) in R(19). We used a difference-in-differences model to identify genes that responded differently between the ablated cirrhotic and ablated non-cirrhotic liver, while controlling for the differences between the unablated samples. The model also accounted for sample pairing (i.e., ablated and control samples taken from the same animal). Over-representation analysis (ORA) and gene set enrichment analysis (GSEA) for gene ontology gene sets were performed with the clusterProfiler (RRID:SCR_016884) package(20). ORA was run on the list of differentially expressed genes (DEGs) with FDR<0.05. GSEA was performed on the list of all genes ranked by log2(fold change). Fold change was calculated using the apeglm method for log fold change shrinkage(21). Gene sets were enriched if FDR<0.05 with Benjamini-Hochberg correction. ORA and GSEA figures were generated with the enrichplot package(22).

### Hyperspectral imaging

Hyperspectral imaging is a technique for measuring the spectra of a pixel over a range of wavelengths by dividing it into many contiguous spectral bands. It can be used with reflectance spectroscopy to non-invasively measure concentrations of oxy- and deoxyhemoglobin and generate a 2D spatial map of tissue oxygen saturation(23).

Hyperspectral images were acquired with the HyperView Imaging System (HyperMed Imaging, Memphis, USA). The camera was held at a fixed distance from the surface of the exposed left lobe, as marked by a laser distance gauge incorporated in the device. Images were acquired immediately before ablation to establish baseline and again at the time of euthanasia. Image acquisition took approximately 1 second. Oxygen saturation maps were generated using the software included on the device.

### Tomato lectin vascular perfusion

Tomato lectin is a common tool for staining perfused vessels. It is most commonly used in mice and delivered via tail vein injection to circulate throughout the body. We found it was not feasible to intravascularly inject enough compound in rats to sufficiently stain the liver. Consequently, we used retrograde perfusion of the left lobe after euthanasia. The hepatic artery and splenic portal vein were clamped shut. 4-0 sutures were used to tie the portal vein shut as well as the branches of the portal vein leading to the medial lobe, right lobe, and caudate lobe. The branch of the hepatic vein coming from the left lobe was cannulated with a 19G needle connected to a short length of silicone tubing, and sutures were tied around the vessel and needle to limit backflow. Fluorescently labelled tomato lectin (Vector Labs, Newark, USA) was diluted to 50µg/mL in PBS immediately prior to injection. 15mL of the tomato lectin solution was injected over approximately 2 minutes. After a further 2 minute incubation period, swabs were gently rolled over the left lobe to help distribute the solution. The portal vein was then cut to allow efflux and another 15mL was injected. Tissue samples were collected for formalin fixation and TUNEL staining.

## Results

### Parameter sweep of fractional laser treatment intensity

As the fractional laser has not previously been used in the liver, we performed a parameter sweep with healthy rats to determine the appropriate treatment settings. The laser variables are pulse energy, number of pulses per microinjury, and density of microinjuries. Our objective was to identify a combination of settings that produced distinct microinjuries, each less than 500µm in diameter and at least 2mm in depth. These results are shown in Fig. S1. Excessive pulse energy or density of microinjuries causes collateral damage to the tissue between ablation sites. Treatment with 15mJx2, 5% density produced microinjuries approximately 300µm in diameter and 2mm deep. This treatment intensity was used for all subsequent experiments.

### Microinjuries are well tolerated in the healthy liver but cause large zones of cell death in the cirrhotic liver

We first investigated the tissue response to fractional laser ablation from 3 days post-ablation up to 14 days. At the 3 day timepoint in the healthy liver, the microinjuries were faintly visible on the surface of the liver (Fig. 1A). Both grossly and histologically, the injuries appeared uniform in size and spacing, and there was no evidence of additional damage beyond the microinjuries (Fig. 1A, C, E). By 7 days, we were no longer able to find any gross or histological evidence of the microinjuries. The cirrhotic liver showed a vastly different response. At 3 days post-ablation, the ablated regions were clearly visible (Fig. 1B). In addition, the injuries were enlarged and distorted. On histology, it was difficult to discern any pattern of the injuries or identify where the individual microinjuries initially were (Fig. 1D, F). There are enlarged and heterogeneous zones of injury that vary greatly in size and shape, with some measuring several millimeters across. HNF4a staining shows there were no hepatocytes within these zones (Fig. 1F). The ablation sites and associated tissue damage were still apparent through 14 days post-ablation. Unablated immunofluorescence reference images are provided in Fig. S2.

**Figure 1:**
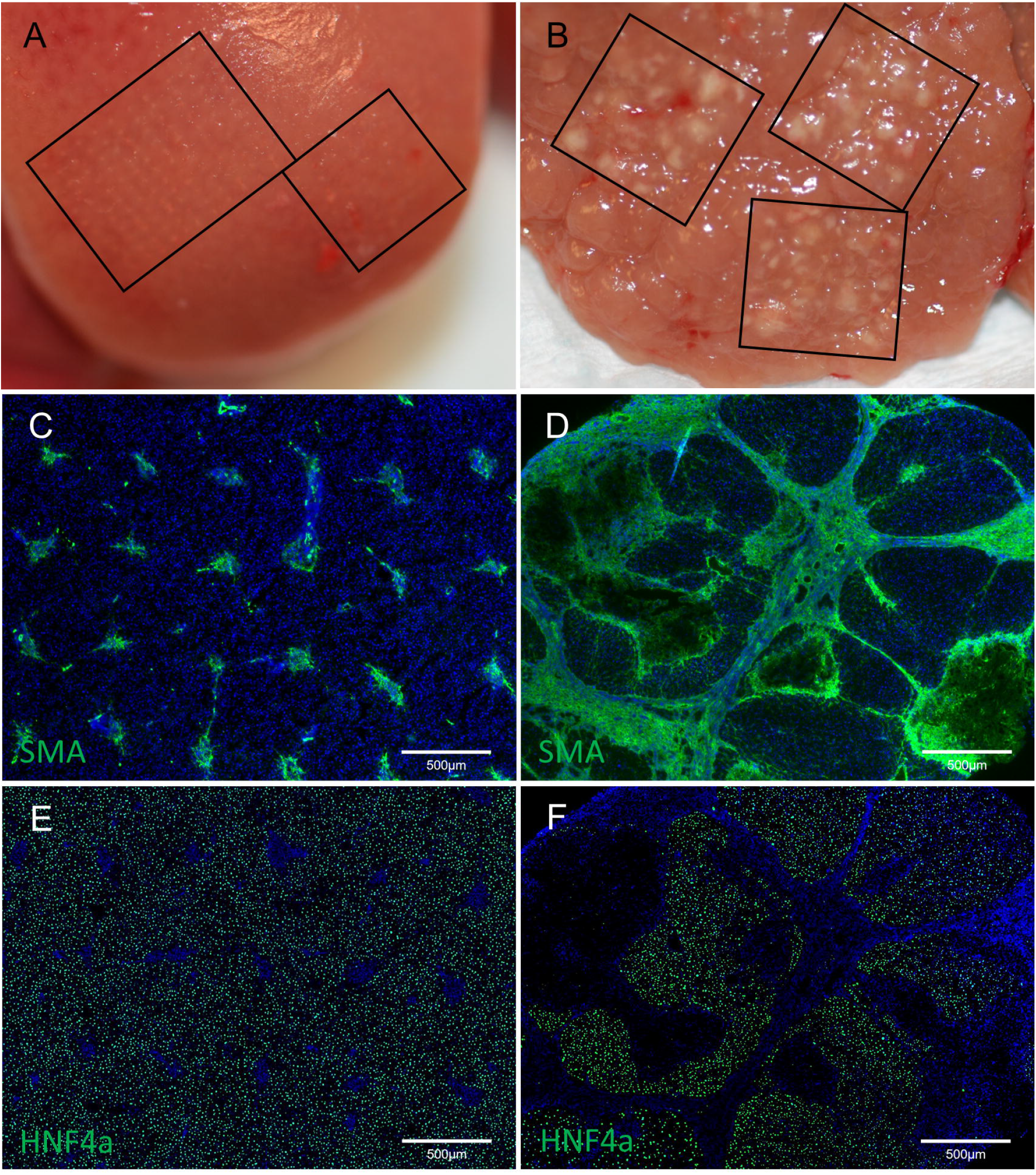
Comparison of response to fractional laser ablation in the healthy (A, C, E) and cirrhotic (B, D, F) liver 3 days after treatment. (A, B) Gross images. Treated regions are outlined with black boxes. The microinjuries in the healthy liver (A) are faintly visible and appear uniform and regular. Injuries in the cirrhotic liver are clearly visible and appear enlarged and misshapen. (C, D) SMA stain showing activated stellate cells recruited to the injuries. Similar to the gross images, the injuries in the healthy liver are small, regular, and uniformly spaced. The injuries in the cirrhotic liver are enlarged and heterogeneous, and the original laser ablation pattern is no longer discernible. (E, F) HNF4a stain for hepatocytes. Hepatocytes are absent from the zones of injury.

To better understand the time course of this response, we looked at earlier timepoints from immediately after ablation up to 6hrs. TUNEL staining immediately after ablation showed a small circle of TUNEL positive cells at each microinjury site where the laser coagulated tissue (Fig. 2A). Little change was observed at 2hrs (Fig. 2B). After 4hrs, cell death was observed beyond the initial ablation sites (Fig. 2C). At 6hrs the enlarged and heterogeneous injury pattern observed at the 3 day timepoint was fully established (Fig. 2D). At this timepoint, the laser injury appears as a bright and dense region of coagulation on TUNEL staining and is easily distinguished from subsequent cell death. These results show that the laser initially produces the expected uniform size and spacing of microinjuries, and the heterogeneous injury zones develop as a response in the hours after ablation.

**Figure 2:**
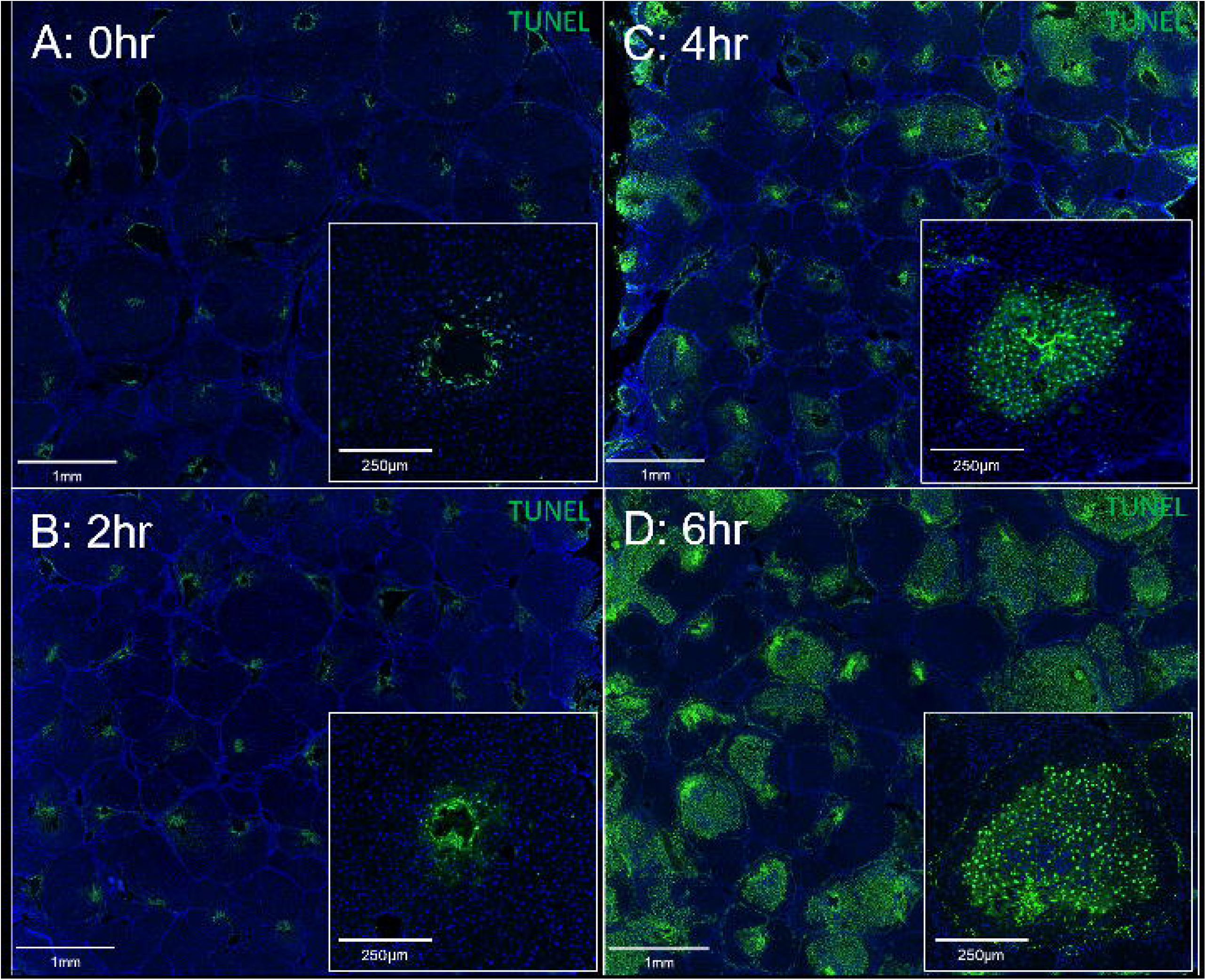
Development of the zones of cell death over the first 6hrs after fractional laser ablation. (A-D) TUNEL stain of the cirrhotic liver at 0hr, 2hr, 4hr, and 6hr after fractional laser ablation, respectively. At 0hr and 2hr, there is little cell death beyond the initial injury from the laser. At 4hr, some zones of cell death begin to develop. The initial injury appears as bright, TUNEL positive coagulated nuclei (cluster of bright green elongated and distorted ribbons). Subsequent cell death is shown by TUNEL positive nuclei with the normal spherical morphology. By 6hr, the exacerbated and heterogeneous injury pattern observed at 3days is established.

### Zones of cell death are heterogenous and influenced by vasculature

There was heterogeneity in the tissue damage pattern, even between adjacent microinjuries. For the majority of injuries, the zone of cell death spreads throughout the cirrhotic nodule containing the microinjury (Fig. 3A). The injury did not spread to adjacent nodules, i.e., there was no cell death in nodules that did not contain a microinjury. This was observed in both small and large nodules. Importantly, we also observed that there was almost always a band of cells next to the septa that survived (Fig. 3A). The band of surviving cells may be more easily seen on the NBTC viability stain (Fig. S3). This was the case even when the microinjury occurred close to the septa and the zone of cell death spread throughout the rest of the nodule (Fig. 3B). Less frequently, we observed injuries that did not spread – even when there were multiple microinjuries within the nodule (Fig. 3C) – or that would only spread out in one direction from the microinjury (Fig. 3D). These patterns of injury were uncommon and were found within the same sections as the more typical injury pattern described above. In the healthy liver, cell death was limited to the region immediately surrounding the laser ablation site, and there was no change from 2 to 6hrs after treatment (Fig. S4).

**Figure 3:**
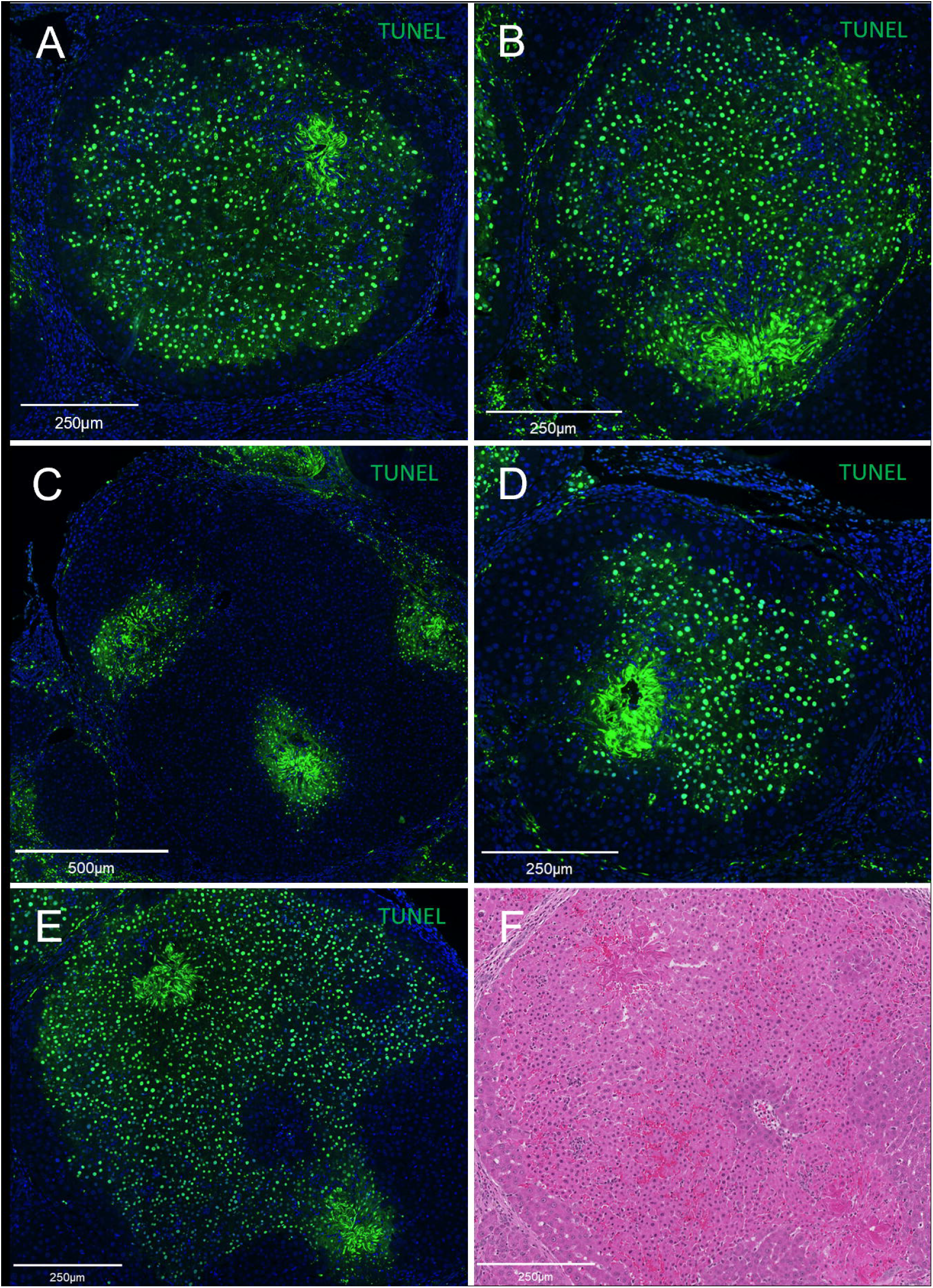
Heterogeneity of response to microinjuries at 6hrs after fractional laser ablation in the cirrhotic liver. (A) The most common nodule injury pattern. The zone of cell death encompasses the entire nodule, except for a band of surviving cells near the septa. (B) Even injuries far to one side of a nodule can cause cell death throughout the majority of the nodule, while cells close to the microinjury but near the septa still survive. (C) Some nodules can contain multiple microinjuries but have very little cell death. (D) Some zones of cell death appear to spread in only one direction from the microinjury. (E) Observation of an island of surviving cells surrounded by TUNEL positive cells. (F) H&E staining of a serial section showed that these surviving cells are near a blood vessel running through the middle of the nodule.

In one section we observed a unique injury pattern of a small island of viable cells surrounded by TUNEL positive cells (Fig. 3E). By comparing this to H&E staining of a serial section (Fig. 3F), we found that the viable cells bordered a larger vessel running through the middle of the nodule – an uncommon anatomical feature at this stage of cirrhosis due to the way the septa form. Although few cases of this were observed, when considered in addition to the observation of the band of viable cells next to the septa – which contain many perinodular vessels – it raised the suspicion that cells are spared by proximity to blood vessels.

### Gene sets related to ischemia, hypoxia, and angiogenesis are enriched after microinjury ablation in the cirrhotic liver

To further characterize the short-term response, we performed RNAseq differential expression analysis with samples from the 4hr timepoint. For our analysis we used a difference-in-differences model to identify genes that responded differently between the ablated cirrhotic and ablated non-cirrhotic liver, while controlling for the differences between the unablated samples. Direct comparison of the ablated cirrhotic and ablated healthy liver would return a list of DEGs dominated by expression differences between healthy and cirrhotic liver unrelated to the microinjury response. The difference-in-differences model allowed us to focus specifically on DEGs underlying the different response to microinjuries between the two tissue states. We identified 520 DEGs with FDR<0.05. ORA of this list of DEGs returned enrichment of gene sets related to ischemia, hypoxia, and angiogenesis, including: response to hypoxia, cellular response to decreased oxygen levels, response to starvation, and regulation of vasculature development (Fig. 4A, B). The DEGs contributing to the enrichment of these gene sets included well-known ischemia related genes such as HIF1a and VEGFA (Fig. 4C). GSEA, which identified enriched gene sets on a ranked list of all genes rather than specifically the DEGs, returned similar results (Fig. S5).

**Figure 4:**
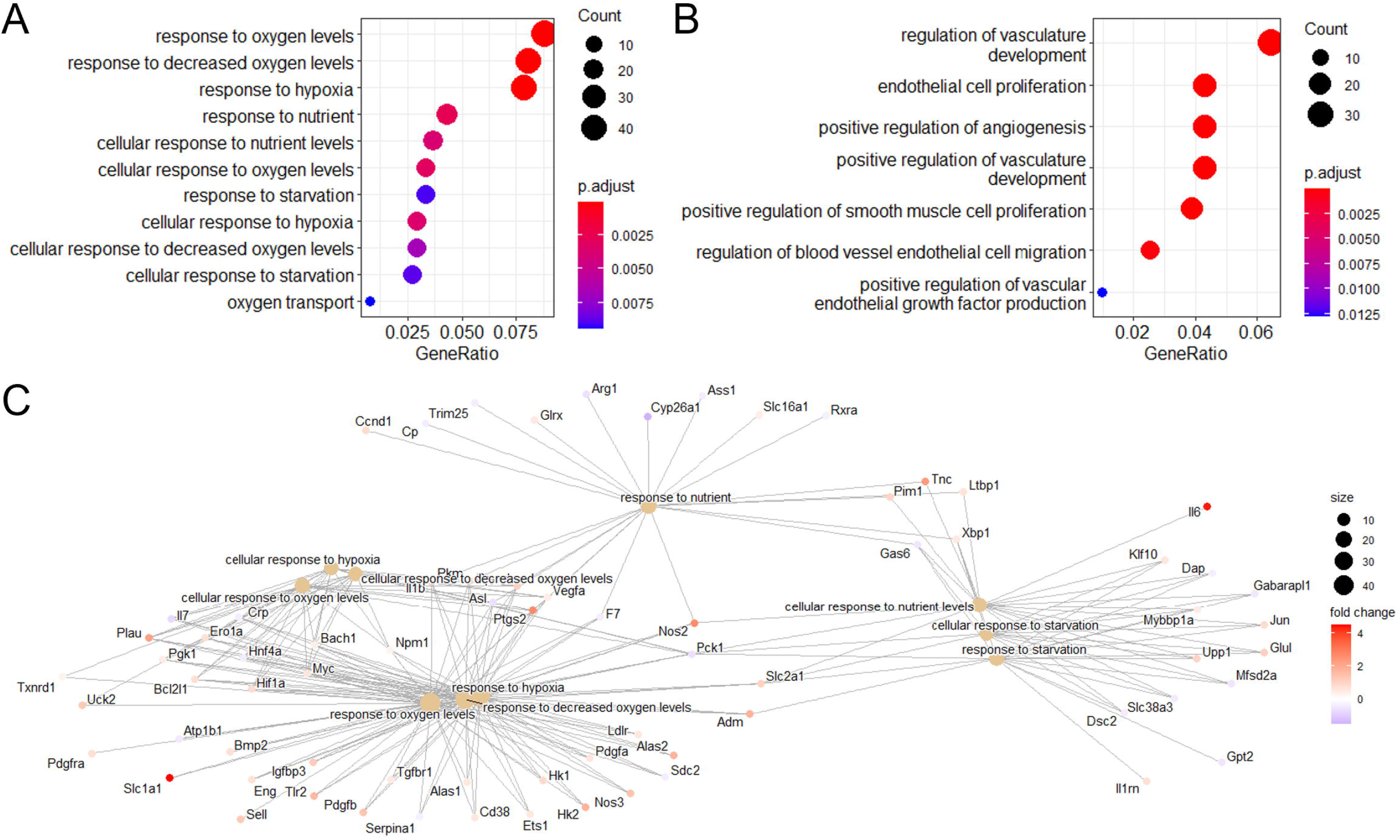
Over-representation analysis (ORA) of RNAseq results comparing healthy and cirrhotic liver at 4hrs after fractional laser ablation. (A) Dotplot of selected enriched gene sets related to ischemia and hypoxia. (B) Dotplot of selected enriched gene sets related to angiogenesis. (C) Concept network plot showing the DEGs contributing to enrichment of select ischemia and hypoxia related gene sets.

### Microinjury ablation reduces tissue oxygen saturation and disrupts perfusion in the cirrhotic liver but not the healthy liver

The RNAseq analysis supported our hypothesis that the exacerbated response to microinjury ablation in the cirrhotic liver was a result of ischemia. To further examine this, we directly investigated the two components of ischemic injury – reduced tissue oxygenation and disrupted blood supply.

Spatial maps of oxygen saturation were acquired with hyperspectral imaging. At the 2hr timepoint in the cirrhotic liver, there was clearly visible decreased tissue oxygen saturation in the areas subjected to microinjury ablation (Fig. 5A). By 6hrs, the ablated regions still had lower oxygen saturation, but it was less apparent (Fig. 5B). This is likely due to rebalancing of supply and demand. We have shown that a large percentage of cells in these areas are undergoing apoptosis at the 6hr timepoint (Fig. 2D), so the oxygen demand would be significantly decreased. There was no apparent decrease in tissue oxygen saturation at 2hrs or 6hrs in the ablated areas of the healthy liver (Fig. 5C, D).

**Figure 5:**
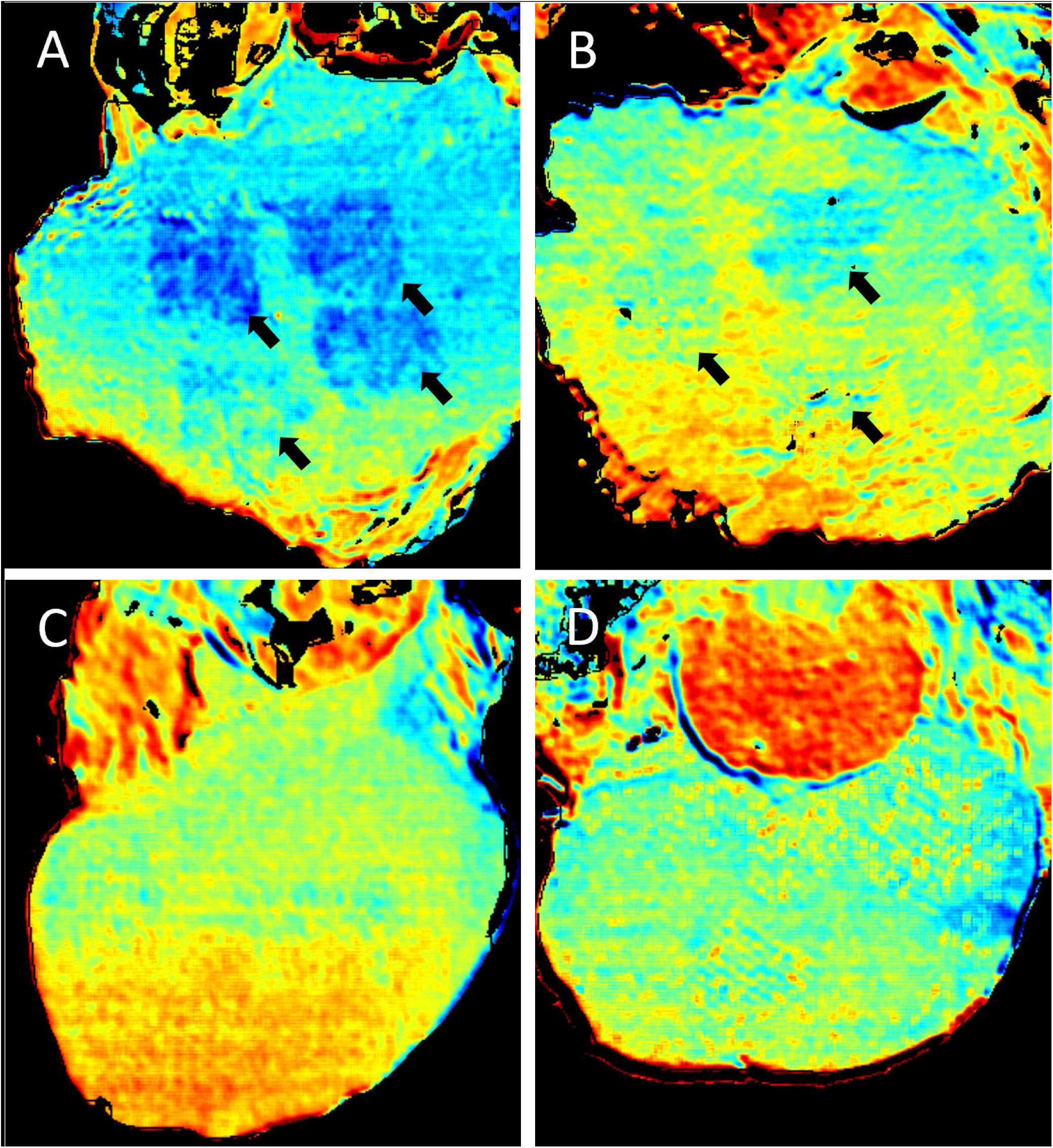
Relative tissue oxygen saturation measured via hyperspectral imaging. (A, B) Cirrhotic liver at (A) 2hrs and (B) 6hrs after fractional laser ablation. At both timepoints, there is clearly reduced oxygen saturation in the regions treated with the laser (indicated with black arrows). This is most distinct at the 2hr timepoint. Healthy liver at (C) 2hrs and (D) 6hrs after fractional laser ablation. Neither timepoint shows a noticeable change in oxygen saturation in the laser treated regions.

The integrity of the blood supply was investigated using tomato lectin perfusion. At the 2hr timepoint in the cirrhotic liver, prior to the development of the zones of cell death on TUNEL staining, large regions of tissue in the ablated areas were not perfused (Fig. 6A; Fig. 7A). We can also identify similar features to the injuries that later develop: many nodules are not perfused at all, some are entirely perfused except around the immediate vicinity of the injury, and some are partially perfused. By 6hrs the tomato lectin stain had the same features as at the 2hr timepoint, but the zones of cell death had now developed on TUNEL staining (Fig. 6B; Fig. 7B). There was no overlap between the tomato lectin and TUNEL staining, i.e., the unperfused regions matched the regions of cell death. The healthy liver maintained perfusion between the individual laser injuries (Fig. 6C, D). An unablated reference image is provided in Fig. S6.

**Figure 6:**
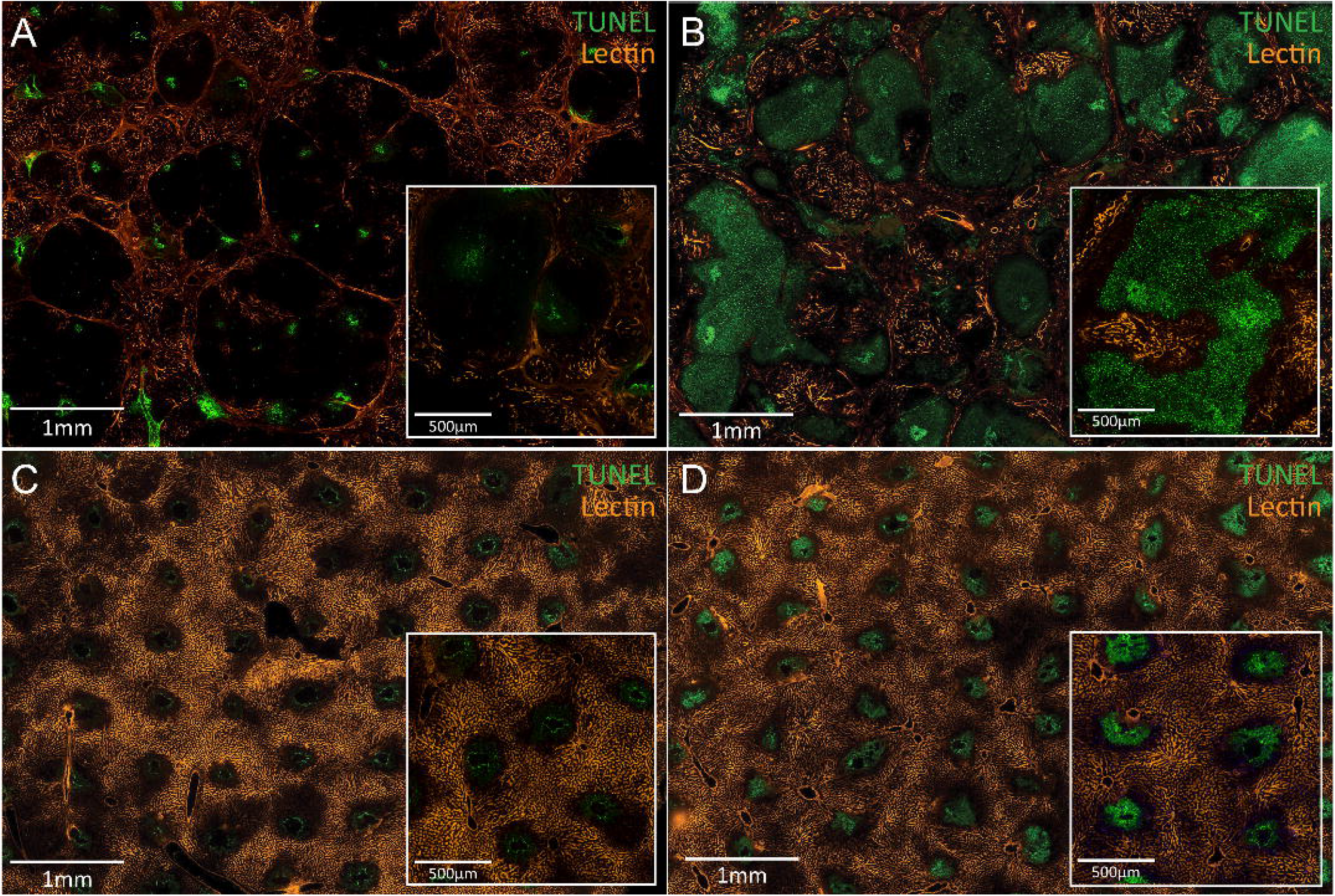
TUNEL and tomato lectin perfusion staining. Cirrhotic liver at (A) 2hrs and (B) 6hrs after fractional laser ablation. At 2hrs, the injuries are still small and uniform, but many nodules are either partially or entirely not perfused. At 6hrs, the enlarged and heterogenous injuries have developed. The injuries align with the unperfused regions of tissue. Healthy liver at (C) 2hrs and (D) 6hrs after fractional laser ablation. At both timepoints, the tissue surrounding the microinjuries remains perfused.

**Figure 7:**
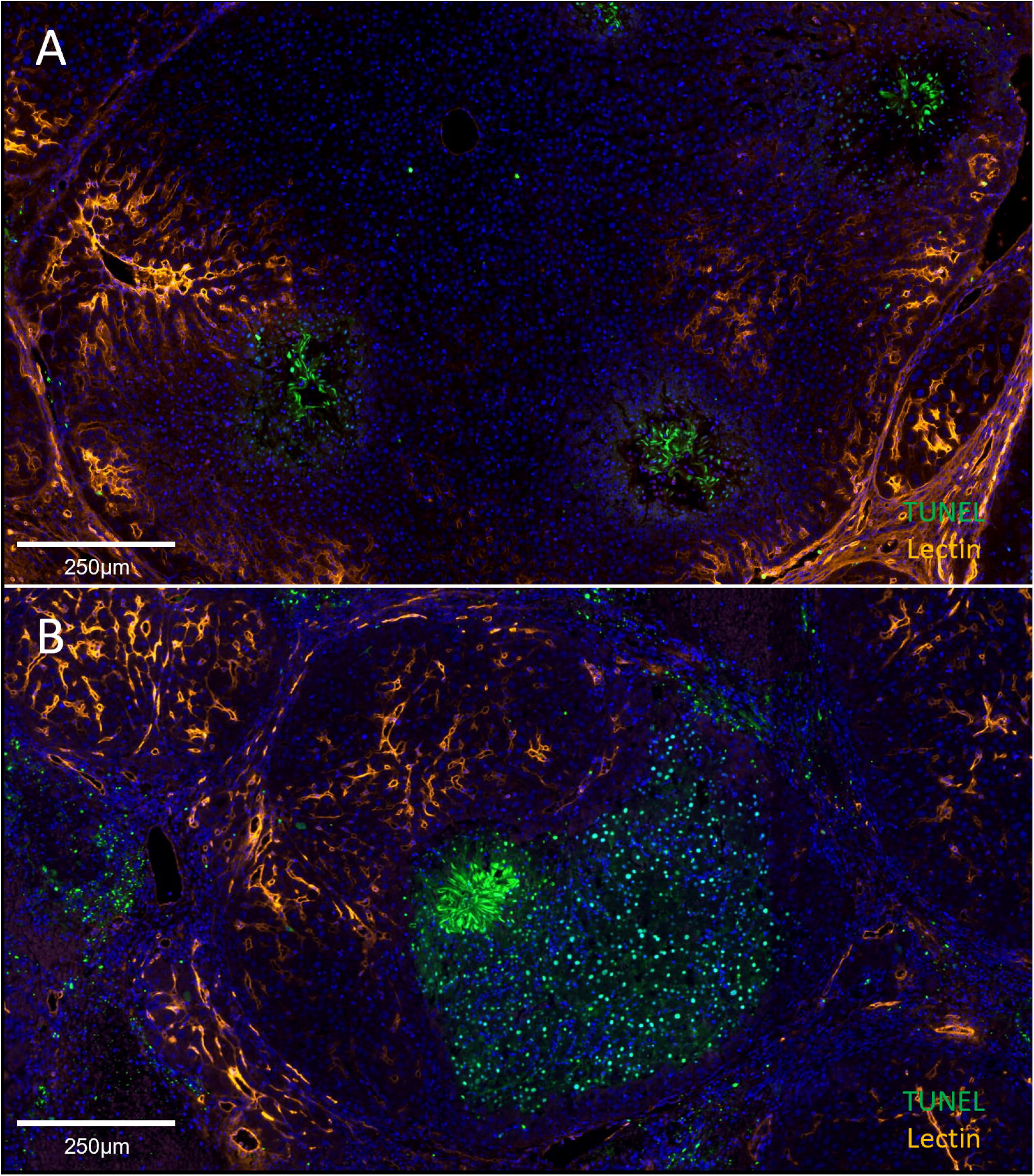
Higher magnification TUNEL and tomato lectin perfusion staining at (A) 2hrs and (B) 6hrs after fractional laser ablation with DAPI counterstain. (A) Small regions of the nodule are perfused, but the majority is not. Cell death is limited to the regions directly ablated by the laser. (B) At 6hrs, the zone of cell death fills the unperfused region of the nodule. The perfused region has no TUNEL positive cells.

A summary of the timeline of the response to microinjuries in the healthy and cirrhotic liver is provided in Fig. 8.

**Figure 8:**
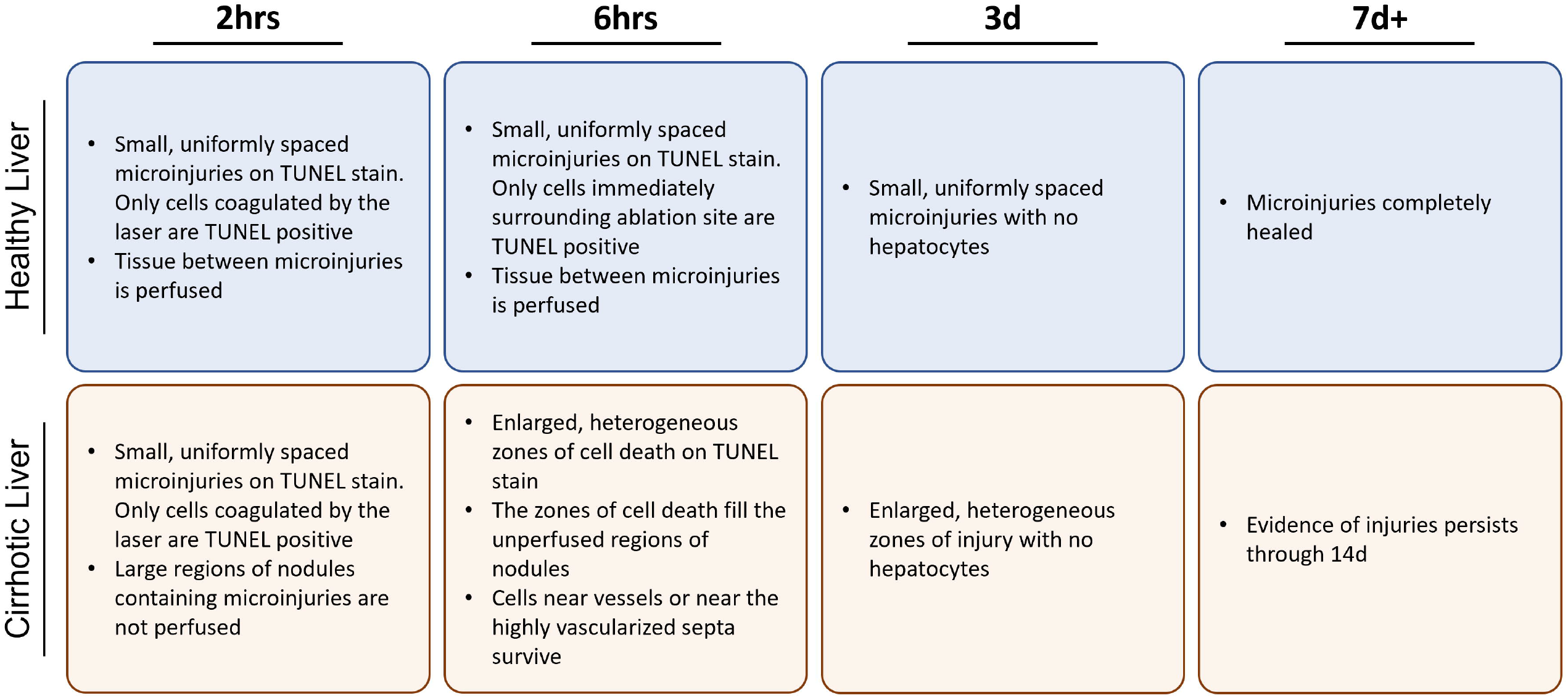
Summary timeline of the response to microinjuries in the healthy and cirrhotic liver

## Discussion

In the skin, the fractional laser enables macrophages and neutrophils to infiltrate the densely packed, highly cross-linked collagen fibrils that make scar tissue resistant to degradation(24, 25). This induces expression of a range of MMPs that cause a potent, short-term matrix remodeling phase(26-28), leading to a significant reduction in scar stiffness within a few days of treatment(29). As a result, the critical positive feedback loop between fibrosis and tissue stiffness is disrupted(25), enabling long-term remodeling towards normal tissue.

In this study, we aimed to characterize the tissue response to fractional laser-induced microinjuries in the healthy and cirrhotic liver to determine whether microinjury ablation can be used to induce scar remodeling in the liver. This technique relies on creating small, finely controlled microinjuries to stimulate remodeling; however, our results showed that the cirrhotic liver instead developed enlarged and heterogeneous injuries. Using RNAseq, hypoxia imaging, and perfusion studies, we have shown that this unexpected pattern of injury is the result of increased susceptibility to ischemia from microvascular damage in the cirrhotic liver.

The liver microvasculature is an essential component of cirrhosis. Liver fibrosis develops through repeated microvascular injuries that cause large regions of hepatocytes to die and be replaced by scar tissue in a process termed parenchymal extinction(30, 31). This progresses over time to the characteristic nodular architecture with the formation of an extensive network of perinodular vessels within the septa(32, 33). These vessels include intrahepatic shunts that allow blood to bypass the parenchyma(34, 35). Capillarization of sinusoids and perisinusoidal fibrosis impair oxygen delivery to hepatocytes, while vascular remodeling and endothelial dysfunction increase intrahepatic resistance and further limit perfusion(36-40). All these factors combine to create a hypoxic environment that makes the cirrhotic liver more susceptible to ischemic injury in the event of reduced blood flow (e.g., ischemic hepatitis(41)). It is important to note that this is distinct from the ischemic susceptibility demonstrated here. Our results show that a nodule is susceptible to an obstruction at the microvascular level (i.e., a laser microinjury) broadly disrupting perfusion.

The ability of a vascular network to maintain perfusion after an obstruction is determined by the extent of collateralization and intra-tree anastomoses(42). These are anastomoses between arterioles that normally have little to no flow across them. Obstruction causes pressure to drop in the affected arteriole, allowing blood to flow through the anastomosis and maintain perfusion(42, 43). These connections can occur between arterial trees (collaterals) or within the same tree (intra-tree anastomoses), though they serve the same role in the event of obstruction(42, 44). In this study, vascular perfusion was maintained post-injury in the healthy liver, indicating sufficient connections to compensate for the damage to the vascular network. In the cirrhotic liver, the changes in the vasculature that occur during the progression of cirrhosis – potentially including changes in collateralization – appear to have significantly reduced the ability to maintain perfusion after a disruption.

In addition, the pattern of injury observed in the cirrhotic liver after laser treatment matches some of the features of the nodule microvasculature. Cells at the edges of the nodule survive even when all sinusoids are not perfused because of the perinodular vasculature, and injuries spreading only within a nodule is consistent with the microvascular network being separate from neighboring nodules(45). The heterogeneity of the injury pattern may reflect differing placement of the microinjury within the microvascular network. If the microinjury hits a critical branching point then perfusion throughout the nodule may be impacted (e.g., Fig. 3A), but if it hits a part of the network with few downstream branches – or where downstream tissue can be fed by a collateral pathway – then the impact would be less severe (e.g., Fig. 3C, D).

Compartment syndrome could also play a role in these ischemic injuries. Compartment syndrome is most commonly seen in muscle, where the parenchyma is divided into compartments by inelastic fascia. Increased pressure within a compartment, e.g. due to hemorrhage, can collapse the microvasculature to prevent perfusion and cause ischemic injury(46). Under this hypothesis, hemorrhage from the microinjury increases the pressure within the compartment formed by the septa. In compartment syndrome, capillaries collapse when the compartment pressure exceeds the perfusion pressure. It is unlikely that either perfusion pressure or compartment pressure to one side of a microinjury would be significantly different from the other, so this explanation alone does not account for cases where perfusion was restricted in one direction extending from the microinjury (e.g., Fig. 3D), which can be more easily explained by a disruption to the microvascular network obstructing flow to downstream branches. However, in some nodules the ischemic injury may result from a combination of direct disruption of the microvascular network and compression of the remaining microvasculature by subsequent hemorrhage. This effect would not be seen in the healthy liver because the parenchyma is not compartmentalized, so excess fluid is more free to flow interstitially. Interestingly, a compartment syndrome-like effect has been proposed as a mechanism for cirrhosis pathogenesis(47). This model involves a feedback loop of congestion, ischemia, and parenchymal extinction, leading to further congestion. It was suggested that the compartmentalization in this case arises from a nested cone angioarchitecture, where each cone is drained by a branch of the hepatic vein, and there is little collateral circulation between cones. However, they speculated that this effect would not be observed in animal models of cirrhosis because of differences in the hepatic venous tree.

Hypoxic injury is an important topic in cirrhosis. The cirrhotic liver is more susceptible to a number of ischemic conditions than the healthy liver, such as ischemia-reperfusion injury during transplantation (48) or ischemic hepatitis from congestive heart failure(41). These types of ischemic injuries are related to pre-existing hypoxia and hepatocellular dysfunction rather than obstruction at the microvascular level. However, there are other ischemic conditions that do involve liver microvascular obstruction, such as sinusoidal obstruction syndrome (SOS). SOS has various causes, including hematopoietic stem cell transplantation and chemotherapy toxicity. It involves damage to the sinusoid endothelium, which causes randomly dispersed plugs of erythrocytes that obstruct blood flow(49). Our findings suggest that SOS would cause much greater tissue damage in the setting of cirrhosis, not only due to pre-existing hypoxia but also because the obstructions would disrupt perfusion to a larger tissue volume. A similar effect would be seen with microinfarctions in the liver, which can be caused by cirrhotic coagulopathy(50).

It is worth noting that the exacerbated injuries were not a result of heat. It is relatively straightforward to mathematically model the temperature change around a microinjury. These calculations and a brief discussion of heat tolerance are provided in the Supplementary Information.

The exacerbated injuries limit the applicability of the current fractional laser tool for inducing scar remodeling in cirrhosis. Alterations to the fractional laser tool could help avoid this issue. Femtosecond lasers can create extremely small injuries, potentially on the size of a single cell. Such a laser would have limited treatment depth, but repetitive pulse stacking with precise targeting may provide sufficient treatment volume. A similar effect could be achieved by focusing the laser to a smaller spot size. A balance would have to be struck between creating small enough injuries to avoid significantly disrupting the microvascular network while still eliciting a sufficiently strong response to drive scar remodeling.

There are numerous fibrotic conditions occurring throughout the body that would not have the same limitations we discovered in the liver. For example, a fractional laser tool combined with an endoscope may be a promising option for the treatment of gastrointestinal strictures – fibrous bands that can progressively contract over time and cause blockages. Another potential use case is pulmonary fibrosis. Similar to cirrhosis, transplantation is the only curative treatment option for pulmonary fibrosis. It may be possible to combine a fractional laser with a bronchoscope to perform minimally invasive microinjury treatment. There is a multitude of fibrotic conditions occurring throughout the body, many of which have high morbidity and limited treatment options. The remarkable results of microinjury treatment in the skin warrant further investigation for other applications.

Our study also introduces a novel controlled method for creating microvascular injuries which can be used to examine functional consequences of the cirrhotic vasculature that would be difficult to obtain through imaging studies or vascular casts alone. Future studies could investigate the response to microinjury at various stages of fibrosis. Shunting vessels have been shown to develop as early as 4 weeks after CCl4 exposure(35), when fibrosis is minimal. Investigating how the injury response to the laser changes over the various liver fibrosis states and at which point the ischemic susceptibility first appears could provide useful information about the time course of vascular derangement. It would also be interesting to investigate the response to fractional laser ablation during the regression of fibrosis. Certain animal models of cirrhosis exhibit reversible fibrosis, even after cirrhosis is established.

However, the vascular changes are not reversible. Studying the response to microinjuries during regression of the septa may provide information about how perfusion changes over time and whether the ischemic susceptibility remains permanently.

We have demonstrated that fractional laser ablation is well tolerated in the healthy liver but causes enlarged and heterogeneous zones of cell death in the cirrhotic liver. These exacerbated injuries result from disrupted perfusion of the cirrhotic nodule. The increased susceptibility to ischemia from microvascular damage is likely related to the vascular derangements that occur during the progression of liver fibrosis to cirrhosis. Modifications to the fractional laser tool, such as using a femtosecond laser or reducing the spot size, may prevent large disruptions of perfusion and enable further development of a laser-induced microinjury treatment for cirrhosis.

## Data Availability

Raw and processed RNAseq data are available at https://www.ncbi.nlm.nih.gov/geo/query/acc.cgi?acc=GSE228374.

## Author Contributions

Conceptualization: BDL, KKT, YVP, JT, RRA. Formal analysis: BDL. Funding acquisition: RRA. Investigation: BDL, MS. Methodology: BDL, MS. Supervision: JT, RRA. Writing – original draft: BDL. Writing – review & editing: MS, KKT, YVP, JT, RRA.

## Acknowledgements

RRA was partially supported by the Lancer Endowed Chair in Dermatology. The graphical abstract was prepared with BioRender.

