## Supplementary Material for "Increased susceptibility to ischemia causes exacerbated response to microinjuries in the cirrhotic liver"

### Supplemental Material

**Table of contents**

Supplementary Information: Thermal Diffusion Calculation and Heat Tolerance Discussion…………. 2

Fig. S1: Injury pattern at 2hrs after fractional laser ablation in the healthy liver for various treatment intensities……………………………………………………………………………………………………………………. 3

Fig. S2: Untreated immunofluorescence reference images for the healthy and cirrhotic liver………………………………………………………………………………………………………………………………………..4

Fig. S3: NBTC viability stain in the cirrhotic liver (A) immediately and (B) 6hrs after fractional laser treatment……………………………………………………………………………………………………………………………….5

Fig. S4: TUNEL stain in the healthy liver (A) 2hr and (B) 6hrs after fractional laser treatment……………………………………………………………………………………………………………………………….6

Fig. S5: Gene Set Enrichment Analysis (GSEA) of RNAseq results comparing healthy and cirrhotic liver at 4hrs after fractional laser ablation…………………………………………………………………………………7

Fig. S6: Tomato lectin reference stain of untreated cirrhotic liver…………………………………………………..8

Supplementary References……………………………………………………………………………………………………. 9

**Supplementary Information: Thermal Diffusion Calculation and Heat Tolerance Discussion**

The equation for thermal diffusion from a pulsed laser is(1):

$$T(r,t) = T_{max}\left(\frac{t_0}{t}\right)^{3/2} exp\left(-\frac{r^2}{4Dt}\right)$$

where $r$ is distance from the center of the laser pulse, $t$ is time, $t_0$ is the laser pulse width, $D$ is thermal diffusivity, and $T_{max}$ is the maximum initial temperature increase induced by the laser. Using this model, the peak temperature increase 100um from the edge of the coagulated tissue is 2.20°C, and at 200um the peak temperature increase is just 0.57°C. A precise threshold for thermal damage is difficult to determine due to the influence of both intensity and duration, varying tissue sensitivities, and heat stress processes activated over a range of temperatures(2). One method for quantifying thermal damage is cumulative equivalent minutes at 43°C (CEM43), where any heat dose is converted to an equivalent number of minutes at a tissue temperature of 43°C(3). 9 CEM43 has been suggested as a general threshold for safe thermal exposure, and the lowest reported CEM43 value for thermal damage in the liver is 9.9(3-5). Given the peak temperature increase 100um from the coagulated tissue is estimated to be just 2.20°C for a duration measured in milliseconds, the bulk heating effect of the laser is negligible.

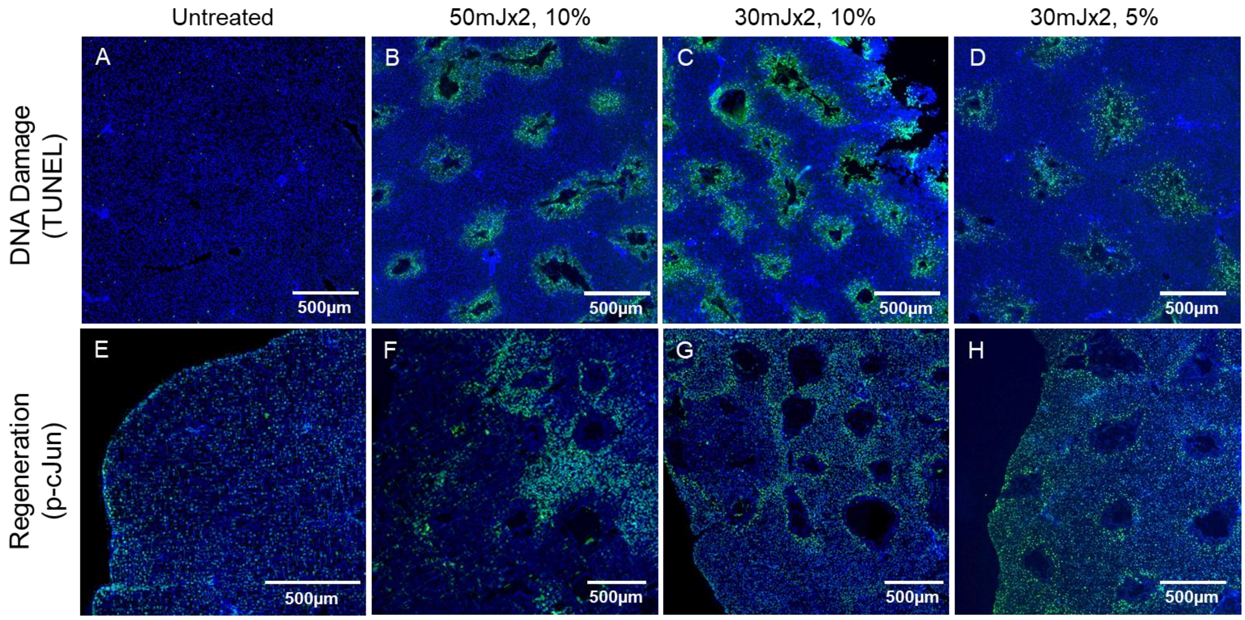


**Fig. S1: Injury pattern at 2hrs after fractional laser ablation in the healthy liver for various treatment intensities.** TUNEL stains DNA damage and shows cells directly injured by the laser. Phospho-cJun is an early marker of liver regeneration and should be expressed in hepatocytes surrounding an injury. The ideal pattern would have positive phospo-cJun staining bordering every individual microinjury (A,E) Untreated tissue has no positive TUNEL staining and homogenous phospho-cJun staining. (B,F) 2 pulses of 50mJ at 10% density. A few injury sites are isolated (clearly surrounded by positive phospho-cJun staining), but most are not, indicating coalescence of the microinjuries into a larger injury. (C,G) 2 pulses of 30mJ at 10% density. Most injuries are isolated but there is one region where several adjacent microinjuries coalesced into a larger injury. (D,H) 2 pulses of 30mJ at 5% density. Reducing the density of injury sites created a clear pattern of isolated microinjuries. 15mJx2, 5% density was then chosen for the remaining experiments to be comfortably below this threshold while still maintaining the desired minimum treatment depth.

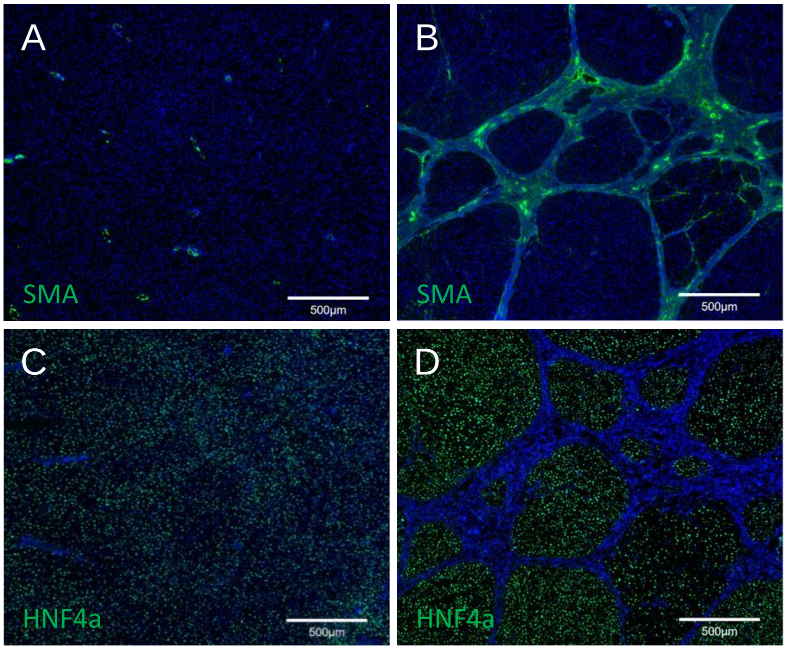


**Fig. S2: Untreated immunofluorescence reference images for the healthy and cirrhotic liver.** (A) SMA stain of healthy liver. (B) SMA stain of cirrhotic liver. (C) HNF4a stain of healthy liver. (D) HNF4a stain of cirrhotic liver.

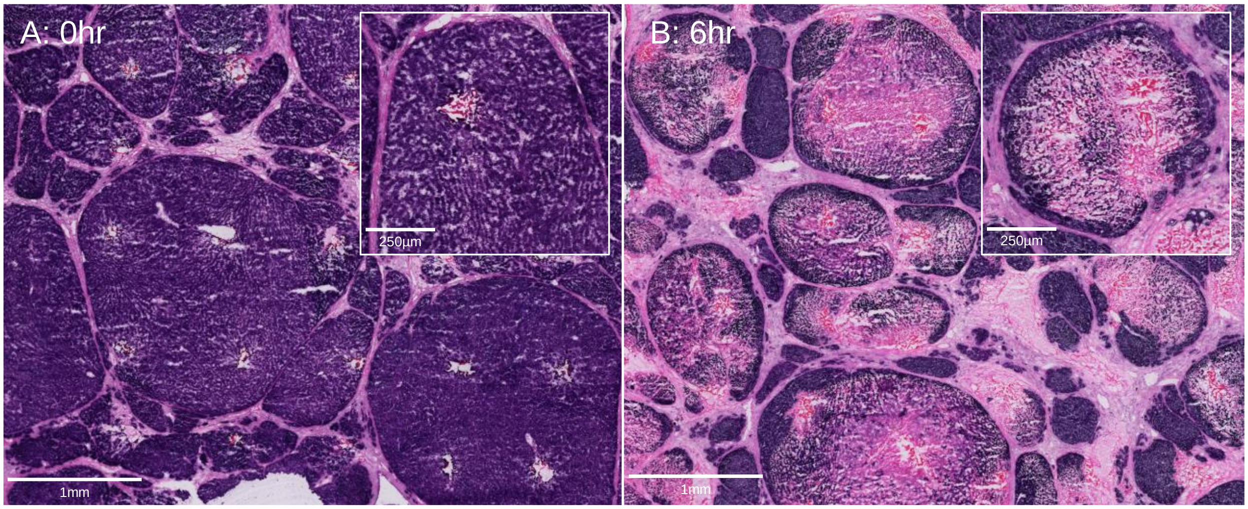


**Fig. S3: NBTC viability stain in the cirrhotic liver (A) immediately and (B) 6hrs after fractional laser treatment.** Viable cells are stained purple. Regions without viable cells are stained pink by the eosin counterstain. (A) Immediately after treatment only the few cells directly ablated by the laser are dead. (B) By 6hr, enlarged and heterogeneous zones of cell death have developed with few viable cells. There is a band of viable cells around the edge of the nodule near the septa.

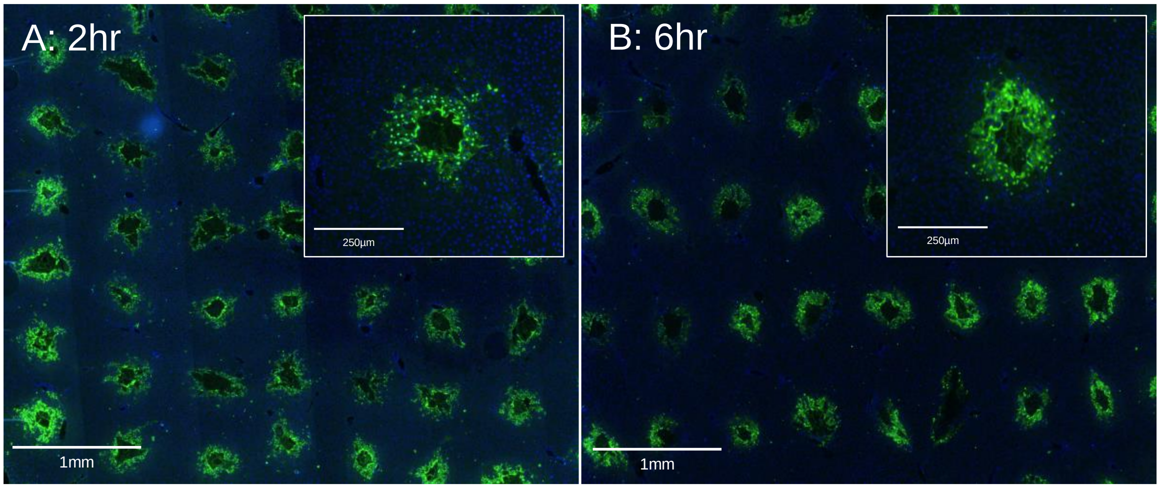


**Fig. S4: TUNEL stain in the healthy liver (A) 2hr and (B) 6hrs after fractional laser treatment.** Cell death is limited to the region immediately surrounding the laser ablation site and there is no change in the injury pattern from 2 to 6hrs.

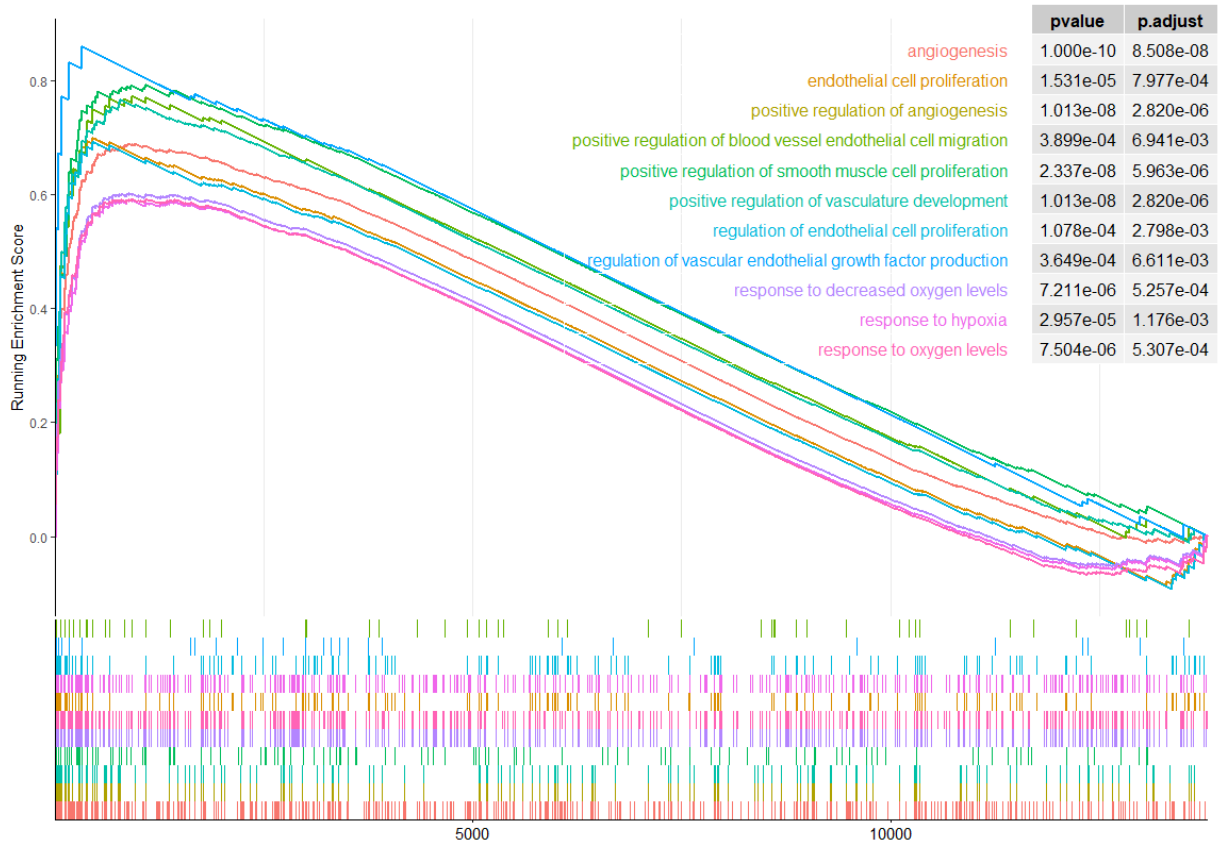


**Fig. S5: Gene Set Enrichment Analysis (GSEA) of RNAseq results comparing healthy and cirrhotic liver at 4hrs after fractional laser ablation.** 4 healthy and 4 cirrhotic animals were used for this analysis, with a laser-treated and untreated control sample taken from each animal. The running enrichment score and adjusted p-value are shown for selected gene sets related to hypoxia and angiogenesis.

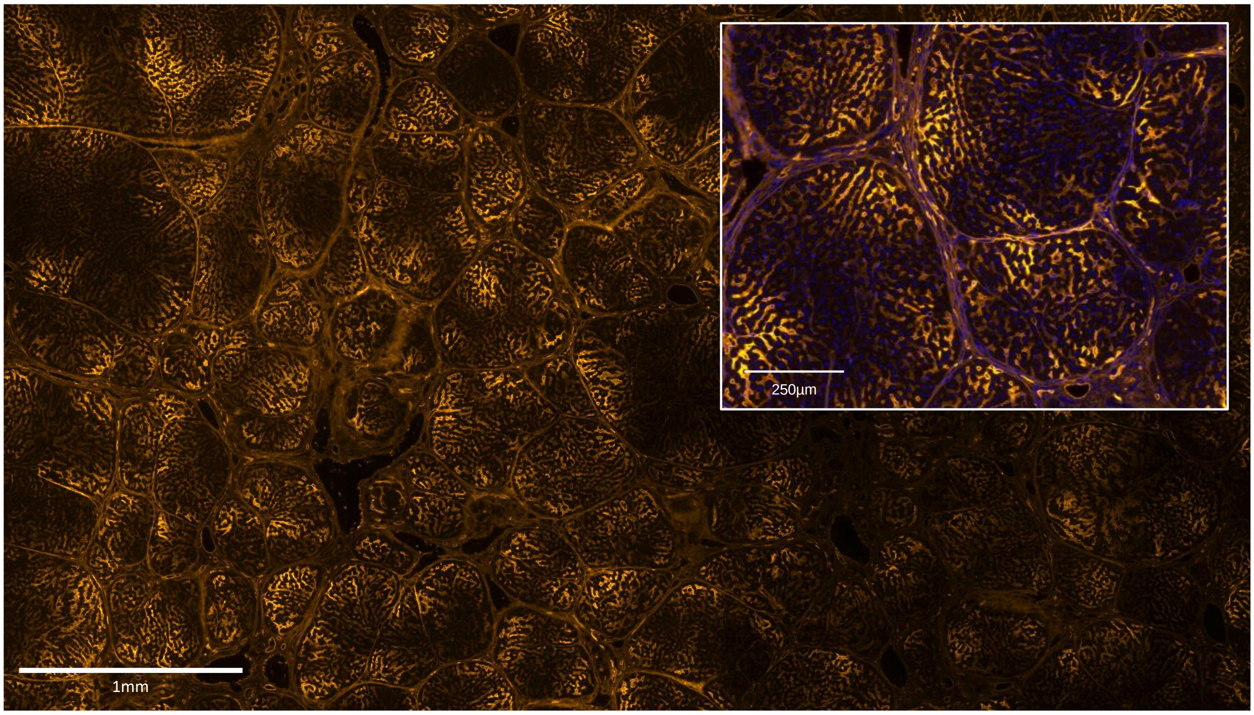


**Fig. S6: Tomato lectin reference stain of untreated cirrhotic liver.** The inset image is also stained with DAPI.
